# Mapping prefrontal afferents along development

**DOI:** 10.1101/2025.09.17.674820

**Authors:** Anne Günther, Malte Stelzer, Andreas Franzelin, Annette Marquardt, Thomas Oertner, Ileana L. Hanganu-Opatz, Jastyn A. Pöpplau

**Author notes:** Corresponding authors: Jastyn A. Pöpplau, Falkenried 94, 20251 Hamburg, Germany, Lead contact, Ileana L. Hanganu-Opatz, Falkenried 94, 20251 Hamburg, Germany. These authors share first authorship (Anne Günther, Malte Stelzer). These authors share last authorship (Ileana L. Hanganu-Opatz, Jastyn A. Pöpplau).

## Abstract

The medial prefrontal cortex (mPFC) acts as a hub for cognitive, emotional, and social processes, integrating neuronal inputs from numerous brain regions. Knowledge on the processes underlying formation and myelination of prefrontal afferents is essential for a comprehensive understanding of the emergence, maturation, and refinement of mPFC-dependent behaviors. However, despite detailed investigation of long-range projections to the mPFC in the adult brain, little is known about their maturation. Here, we use retrograde labeling, light-sheet imaging, and automatic cell segmentation to quantify afferent projections to the mPFC from neonatal to adult age. We show that densities of ipsi- and contralateral prefrontal afferents change along development from a widespread bilateral distribution at neonatal age to a predominantly ipsilateral organization. Furthermore, myelination of interhemispheric prefrontal afferents starts only after the decline of neonatal contralateral projections and is followed by temporally elevated projection densities from limbic brain regions. Overall, these distinct developmental dynamics of prefrontal afferents might have major implications for the maturation of mPFC-dependent functional and behavioral outputs.

## 1. Introduction

The mPFC, a central hub of integrative processing in the brain, is involved in a wide range of cognitive, emotional, and social functions. It integrates direct axonal inputs from numerous cortical and subcortical regions through local and long-range circuits to guide behavior (Hoover and Vertes, 2007; Ährlund-Richter et al., 2019; Hanganu-Opatz et al., 2023). Within the PFC, functions are anatomically and physiologically compartmentalized, with medial subdivisions (*i.e*., anterior cingulate (ACA), prelimbic (PL), and infralimbic (ILA) areas) primarily contributing to sensorimotor functions, action-planning, and decision making, while ventral regions (*i.e*., orbitofrontal cortex (ORB)) as well as adjacent cortices (*i.e*., secondary motor cortices (MOs)) control emotional and motivational behaviors, respectively (Heidbreder and Groenewegen, 2003; Laubach et al., 2018; Anastasiades and Carter, 2021). In contrast, long-range inputs to the mPFC provide essential information on environmental context and internal state from other regions (Spellman et al., 2015; Burgos-Robles et al., 2017). For instance, inputs from the ventromedial thalamus (VM) to the mPFC regulate arousal states (Honjoh et al., 2018), while hippocampal projections contribute to working, recognition, and episodic memory (MacDonald et al., 2011; Takehara-Nishiuchi, 2014; Chao et al., 2017, 2020).

However, not only ipsilateral projections shape prefrontal circuits. Interhemispheric connectivity has been shown to increase the complexity of prefrontal networks (Floresco and Ghods-Sharifi, 2007; Chao et al., 2016; Wang et al., 2023). Numerous studies have investigated the projections crossing the corpus callosum (CC) in rodents and humans (Yorke Jr. and Caviness Jr., 1975; van der Knaap and van der Ham, 2011; Innocenti et al., 2022). While callosal connectivity has been traditionally assumed to be predominantly homotopic (*i.e*., mPFC to mPFC) (Yorke Jr. and Caviness Jr., 1975; De Benedictis et al., 2016), more recent studies have revealed substantial heterotopy of these projections (Szczupak et al., 2023). Notably, specific functions of homotopic and heterotopic callosal connections have been described in rodents. On the homotopic level, sensory processing has been shown to be influenced by contra- and ipsilateral ACA projections (Wang et al., 2023), whereas disruptions of heterotopic pathways between basolateral amygdala (BLA) and ACA lead to impaired decision making (Floresco and Ghods-Sharifi, 2007).

In contrast to this detailed dissection of adult prefrontal afferents, little is known about their development. Projections from other brain regions reach the mPFC during development and are thought to undergo dynamic reorganization during early life to promote maturation of higher-order cognitive (*e.g*., working memory) and emotional functions (*e.g*., fear learning) (Chini and Hanganu-Opatz, 2021). For example, projections from the mediodorsal thalamus (MD) and from the BLA progressively increase in density during development and modulate the maturation of local prefrontal circuits (Cunningham et al., 2002; Maren and Quirk, 2004; Rios and Villalobos, 2004; Parnaudeau et al., 2018; Klune et al., 2021). However, prior studies along development have focused on isolated pathways and time points, precluding a large-scale age-dependent mapping of afferents.

In this study, we filled this knowledge gap and systematically monitored the development of axonal projections to the mPFC from birth to adulthood. Using retrograde viral tracing in combination with light-sheet microscopy, we generated a comprehensive atlas of distinct regional afferent projection patterns to the mPFC during early development. Based on these results, we used the same retrograde viral tracing approach combined with confocal microscopy to apply automatic cell segmentation for quantification of projecting cells from different regions of the brain along the entire development. Additionally, we assessed the degree of myelination of retrogradely labeled inputs using immunohistochemical staining of myelin. Our data provide a comprehensive, brain-wide (*i.e*., ACA, PL, IL, ORB, MOs, agranular insular cortex (AI), piriform cortex (PIRI), BLA, basomedial amygdala (BMA), retrosplenial cortex (RSP), entorhinal cortex (ENT), ventral and intermediate CA1 (CA1v, CA1i), ventral and dorsal subiculum (SUBv, SUBd), claustrum (CLA), anterior nuclei (ATN), VM, MD, supramammillary nucleus (SUM), periaqueductal gray (PAG), and ventral tegmental area (VTA)) developmental overview, revealing distinct, region- and age-specific dynamics of prefrontal afferent and giving crucial insights into the evolving architecture of prefrontal circuits.

## 2. STAR Methods

### 2.1. Key resource table

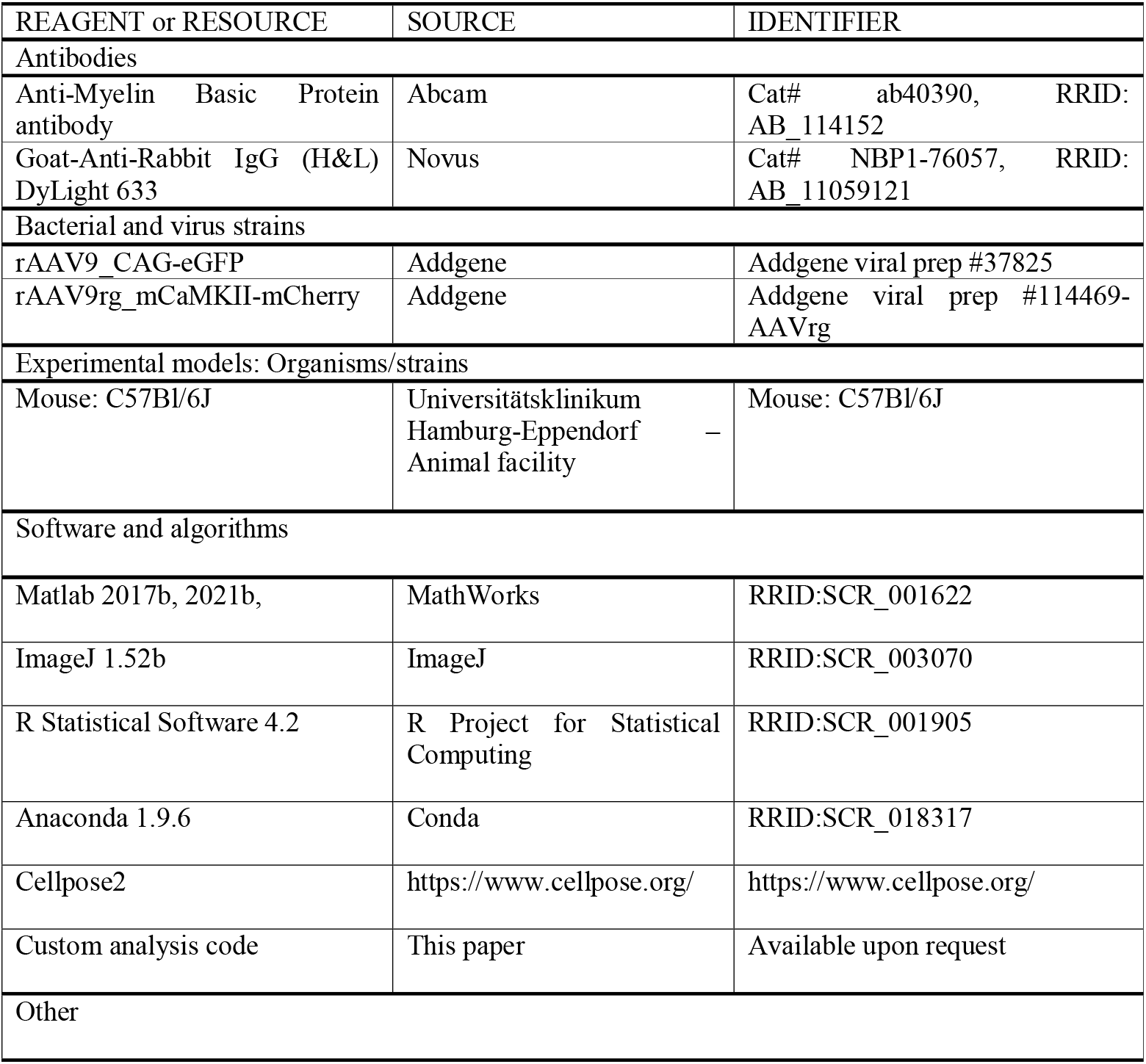
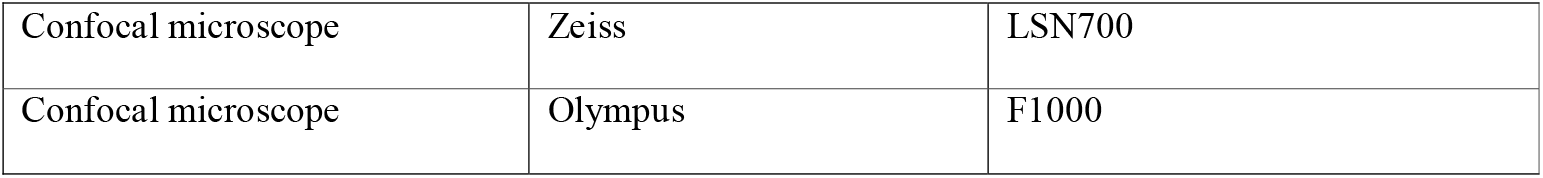

| REAGENT or RESOURCE | SOURCE | IDENTIFIER |
| --- | --- | --- |
| Antibodies |  |  |
| Anti-Myelin Basic Protein antibody | Abcam | Cat# ab40390, RRID: AB_114152 |
| Goat-Anti-Rabbit IgG (H&L) DyLight 633 | Novus | Cat# NBP1-76057, RRID: AB_11059121 |
| Bacterial and virus strains |  |  |
| rAAV9_CAG-eGFP | Addgene | Addgene viral prep #37825 |
| rAAV9rg_mCaMKII-mCherry | Addgene | Addgene viral prep #114469-AAVrg |
| Experimental models: Organisms/strains |  |  |
| Mouse: C57Bl/6J | Universitätsklinikum Hamburg-Eppendorf Animal facility | Mouse: C57Bl/6J |
| Software and algorithms |  |  |
| Matlab 2017b, 2021b, | MathWorks | RRID:SCR_001622 |
| ImageJ 1.52b | ImageJ | RRID:SCR_003070 |
| R Statistical Software 4.2 | R Project for Statistical Computing | RRID:SCR_001905 |
| Anaconda 1.9.6 | Conda | RRID:SCR_018317 |
| Cellpose2 | https://www.cellpose.org/ | https://www.cellpose.org/ |
| Custom analysis code | This paper | Available upon request |
| Other |  |  |
| Confocal microscope | Zeiss | LSN700 |
| Confocal microscope | Olympus | F1000 |

### 2.2. Animals

All experiments were performed in compliance with German law and guidelines of the European Union for the use of animals in research (European Union Directive 2010/63/EU) and were approved by the local ethical committee (Behörde für Gesundheit und Verbraucherschutz Hamburg, N19/121, N23/110). Experiments were carried out in C57BL/6J mice of both sexes. Animals were housed in a 12 h light/12 h dark cycle and were given access to water and food *ad libitum*. The day of birth was considered postnatal day (P) 0.

### 2.3. Virus injections

Viral vectors were injected unilaterally into the PFC of the right hemisphere of C57BL/6J mice. Two viral vectors were applied simultaneously: rAAV9rg_mCaMKII-mCherry (#114469-AAVrg, Addgene, Watertown, MA, United States) for retrograde labeling of excitatory projection-neurons and rAAV9_CAG-eGFP (#37825 Addgene, Watertown, MA, United States) for local labeling of cells at the injection site. Viral vectors were prepared in a genome copy ratio of 1:2.5 and injected at the same time using a stereotax (Kostka and Bitzenhofer, 2022) (Figure 1A). Viral vectors were injected at four different time points, for which volumes and coordinates were adjusted to compensate for brain volume to result in a comparable virus spread across development, *i.e*. P0 (neonatal, 130 nl at a rate of 120 nl/min, 0.3 mm posterior to the inferior cerebral vein, 0.2 mm lateral to the midline and 0.4-0.8 mm ventral from the brain surface), P10 (pre-juvenile, 170 nl at a rate of 120 nl/min, 1,6 mm anterior to Bregma, 0.2 mm lateral to midline and 0,8-1,2 mm ventral from the brain surface), P20 (juvenile, 190 nl at a rate of 120 nl/min, 1,8 mm anterior to Bregma, 0.2 mm lateral to midline and 1-1,4 mm ventral from the brain surface), and P50 (adult, 210 nl at a rate of 120 nl/min, 1,9 mm anterior to Bregma, 0.2 mm lateral to midline and 1,1-1,5 mm ventral from the brain surface). Mice were transcardially perfused with 4% paraformaldehyde (PFA) 10 days after virus injection, *i.e*. at P10 (neonatal, n=6), P20 (pre-juvenile, n=6), P30 (juvenile, n=6), and P60 (adult, n=6). To further account for differences of viral spread at the injection site, brains were manually categorized as low spread, normal spread, or high spread. Low spread was defined as expression in only one mPFC subdivision or a very low degree of labeled cell bodies in more than one subdivision (neonatal n=1, pre-juvenile n=1). High spread was defined as labeled cell bodies in all three subdivisions of the mPFC, while also slightly exceeding mPFC boundaries on the dorsal-ventral or anterior-posterior axis (neonatal n=1, pre-juvenile n=1, juvenile n=1, adult n=1). Accordingly, normal spread was defined as expression in at least two subdivisions without exceeding mPFC boundaries (neonatal n=4, pre-juvenile n=4, juvenile n=5, adult n=5). Virus spread was included in statistical analysis (see Statistical analysis, Supplementary discussion, and Statistics table S1&2).

**Figure 1.**
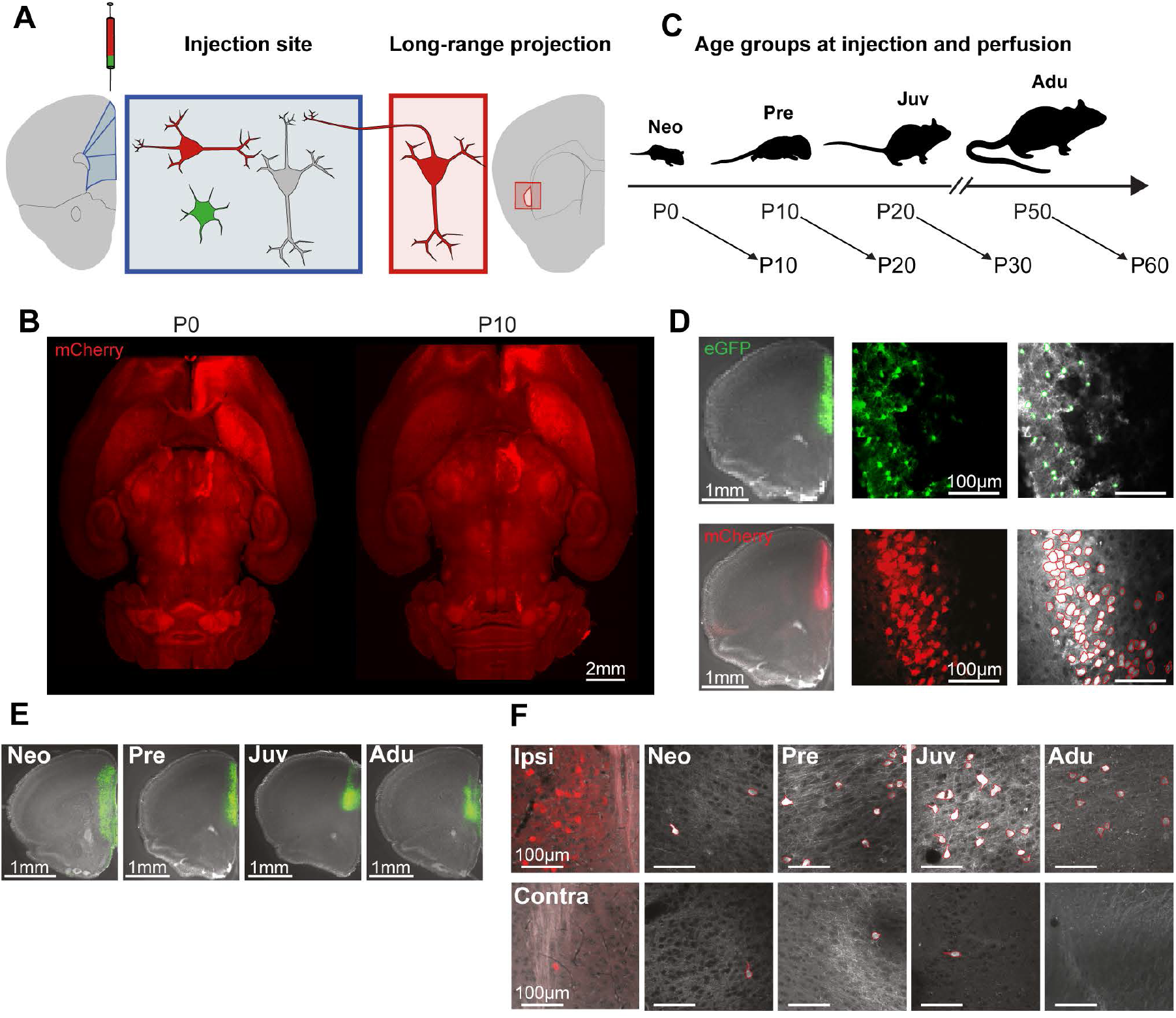

### 2.4. Tissue clearing and light-sheet imaging

Light-sheet imaging on an open source mesoSPIM (Voigt et al., 2019) was used to generate brain-wide overviews of mPFC-projecting brain regions at neonatal (injected at P0, perfused at P10) and pre-juvenile (injected at P10, perfused at P20) age. Perfusion-fixed brains were cleared with SHIELD reagent (Park et al., 2019) to preserve endogenous fluorescence and imaged in EasyIndex solution (RI = 1.465, Life Canvas Technologies) with an Orca Flash 4.0 camera (Hamamatsu).

### 2.5. Cell quantification

For confocal microscopy, brains were prepared in coronal 100 µm sections and mounted on slides using Vectashield with DAPI (Vector Laboratories, Burlingame, CA, United States). For cell quantification, images were obtained using a confocal microscope (LSN700, Zeiss, Germany) with a 20x objective (2,048 × 2,048 pixels; 319.5 × 319.5 µm). Areas were demarcated according to the coronal P56 mouse reference atlas from the Allen Brain Atlas (mouse.brain-map.org). Additionally, images of mice injected at P0 were also compared to a developing mouse brain atlas (Paxinos et al. 2007). Images of three consecutive coronal brain slices were taken for each brain region in the ipsi- and the contralateral hemispheres. Images were segmented with Cellpose 2 in Anaconda (Python 3.8.) using both the standard model algorithm ‘cyto2’ for the images of cells transduced by rAAV9rg_CaMKII-mCherry and a self-trained algorithm as described for the human-in-the-loop approach (Pachitariu and Stringer, 2022) for rAAV9_CAG-eGFP-transduced cells. After automatic cell segmentation, all images were validated by visual inspection. Confocal images from 22 prefrontal and long-range projecting brain sections with mCherry-labeled pyramidal neurons were quantified as a measure of projection strength to the mPFC across development. Images of eGFP-labeled cells were acquired from prefrontal areas at the injection site (ACA, PL, ILA, ORB, MOs) for normalization of quantification of retrogradely labeled projections. Normalized cell count (NCC) was calculated per brain area using mCherry-labeled cells of a single image from each brain area of an animal (r) and all eGFP-labeled cells of an animal (g□) as: 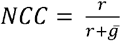 Additionally, the mean cell count (x) was calculated as the mean of mCherry-labeled cells from all sections of one region for all animals from all age groups.

### 2.6. Immunohistochemistry

Immunohistochemical staining of myelin basic protein (MBP) at the CC was performed on the corresponding sections of injected brains after perfusion at P10 (n=4), P20 (n=3), P30 (n=3), and P60 (n=3) as previously described (Parbst et al., 2025). Briefly, brain slices were permeabilized using 0.8% Triton X-100 (Sigma-Aldrich, MO, USA) and blocked with 5% normal goat serum and 5% normal donkey serum (Jackson Immuno Research, PA, USA) for 1 h. Samples were incubated overnight at 4°C with an MBP antibody (1:500, Abcam ab40390). As a secondary antibody Goat-Anti-Rabbit IgG (H&L) DyLight 633 (1:1000, Novus, NBP1-76057) was used.

### 2.7. Myelin quantification

For myelin quantification, images were obtained using a confocal microscope (F1000, Olympus, Tokyo, Japan) with a 60x objective and 2x zoom (1024 × 1024 pixels; 105.47 × 105.47 mm). Immunohistochemically stained samples from injected brains were used to image axonal fibers and myelin at the mid-section of the CC. For every animal, 3-4 stacks from different brain slices (10-30 single images, 150 nm interval) were acquired. After preprocessing and thresholding in ImageJ, colocalization of MBP-positive and mCherry-labeled structures expression was calculated in Matlab R2023a using a custom-written algorithm similar to (Pöpplau et al., 2024). Values were normalized for stack depth.

### 2.8. Statistical analysis

Statistical analysis was carried out in R using linear mixed effect models (LME) of the ‘lme4’ package in R 4.4.1. We assessed the impact of *age* (fixed effect) on NCCs, while accounting for random effects, including *sex, viral spread*, and *image number* to consider for multiple image-based measurements per *individual animal*. Analyzed image stacks of myelinated structures, labeled axonal fibers, and their colocalization were similarly analyzed with LME models to determine the effect of fixed and random effects on myelinization of axons. Briefly, likelihood ratio tests were performed to compare log-likelihoods between the full and the reduced model. To test for a significant effect of age-group (fixed effect), sex, or viral spread (random effects), these parameters were removed from the full model to calculate statistical significance using the ‘anova’ function. Post-hoc pairwise comparisons between age-groups were conducted using the Tukey’s Honest Significant Difference test using the ‘multcomp’ package. P-values were adjusted using the Holm method ‘holm’ to account for multiple comparisons (see Statistics table S1).

## 3. Results

### 3.1. mPFC afferents show age-dependent trajectories

Resolving the developmental dynamics of long-range projections to the mPFC from different brain regions and between hemispheres is the basis for understanding how prefrontal circuits gain and develop their functions. In a first step, we assessed the maturation of prefrontal afferents during early development using light-sheet microscopy on cleared brain tissue. For this, we injected rAAV9rg_mCaMKII-mCherry and rAAV9_CAG-eGFP unilaterally into the mPFC at two different time points (*i.e*., neonatal P0 and pre-juvenile P10). Tissue was perfused and cleared 10 days after injection (*i.e*., neonatal P10, pre-juvenile P20) (Figure 1A, B). Whole-brain imaging revealed retrogradely labeled cells in multiple cortical and subcortical brain regions of both hemispheres at both investigated time points (Figure 1B). Based on this qualitative overview, we chose 22 brain regions with direct axonal projections to the mPFC along development to analyze in detail. We also included the PAG and VTA (Figure 1B), that showed prominent glutamatergic afferent projections during development which is not reported for the adult brain (Hoover and Vertes, 2007; Ährlund-Richter et al., 2019). Next, we used confocal microscopy in slices of injected brains across the entire development (*i.e*., neonatal P0, pre-juvenile P10, juvenile P20, adult P50). Subsequent cell segmentation (Figure 1C-F) to assess projection strength by calculating the NCC (normalized cell count: calculated as ratio of mCherry-labeled cells from each brain area *vs*. all eGFP- and mCherry-labeled cells, see Materials and Methods section) in prefrontal subdivisions (ACA, PL, ILA, ORB, MOs) as well as in cortical (AI, PIRI, BLA, BMA, RSP, ENT, CA1v, CA1i, SUBv, SUBd) and subcortical areas (CLA, ATN, VM, MD, SUM, PAG, VTA), projecting ipsi- and contralaterally to the mPFC.

### 3.2. Medial but not adjacent local prefrontal connectivity increases along development

In order to assess local prefrontal connectivity, we quantified retrogradely labeled cells in prefrontal subdivisions of the ipsilateral hemisphere (marked as ^i^ for all regions). The density of labeled cells in mPFC and adjacent subdivisions changed with age but was not influenced by sex or viral spread (Figure 2A-E). For the ACA^i^ and PL^i^ the NCC increased from neonatal to pre-juvenile age and remained stable afterwards (Figure 2A,B). Solely ILA^i^ local connectivity peaked at juvenile age and decreased in adults (Figure 2C). In contrast, the NCC in adjacent MOs^i^ and ORB^i^ did not significantly change with age (Figure 2D,E). Taken together, these data indicate that especially PL and ILA become strongly integrated into local prefrontal connectivity during development.

**Figure 2.**
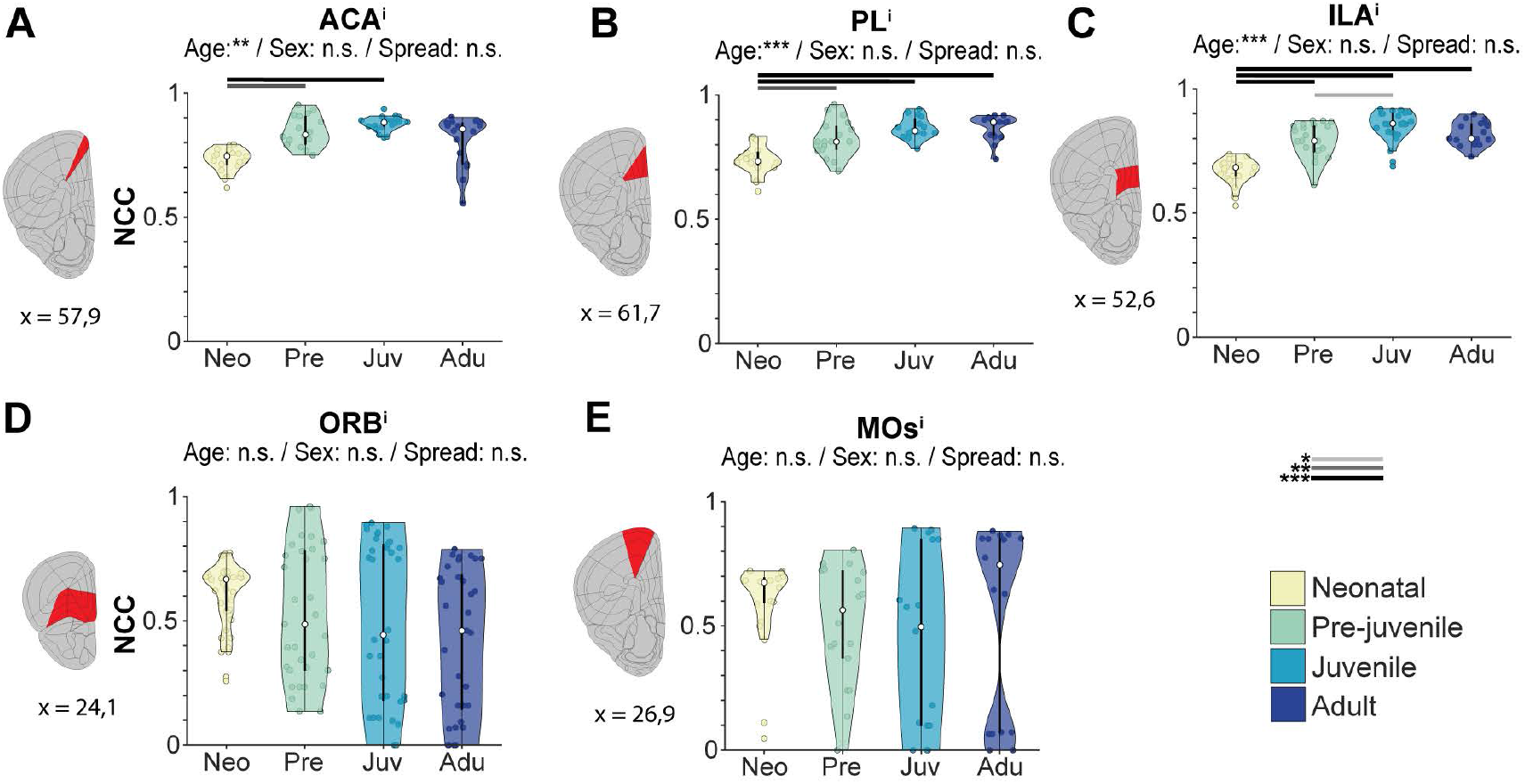

### 3.3. Cortical long-range afferents to the mPFC peak at (pre-)juvenile age

Next, we quantified the densities of long-range afferents from ipsilateral cortical (RSP^i^, PIRI^i^, AI^i^, ENT^i^, SUBv^i^, SUBd^i^, CA1i^i^, and CA1v^i^) as well as cortical adjacent regions (BLA^i^, BMA^i^, CLA^i^). Sex of investigated mice or viral spread had no confounding effects on the results. In most regions, with exception of AI^i^ that showed a consistently high density of afferents, and CA1i^i^ (Figure 3A,B), prefrontal afferents revealed distinct age-dependent patterns. The densities of projections from BMA^i^, CLA^i^, and BLA^i^ strongly increased from neonatal to pre-juvenile age and showed only slight changes later on (Figure 3C-E). Notably, BLA^i^- to-mPFC afferents were nearly absent at neonatal age, emerging towards pre-juvenile age. A similar pattern, though less striking, was also detected for RSP^i^, ENT^i^, and SUBv^i^ (Figure 3F-I,K). However, the densities of these prefrontal afferents decreased during later development, resulting in similar values when comparing neonatal and adult mice. A rather unusual developmental dynamic was detected for SUBd^i^ with dense projections to the mPFC at neonatal age that almost disappeared at adult age (Figure 3J).

**Figure 3.**
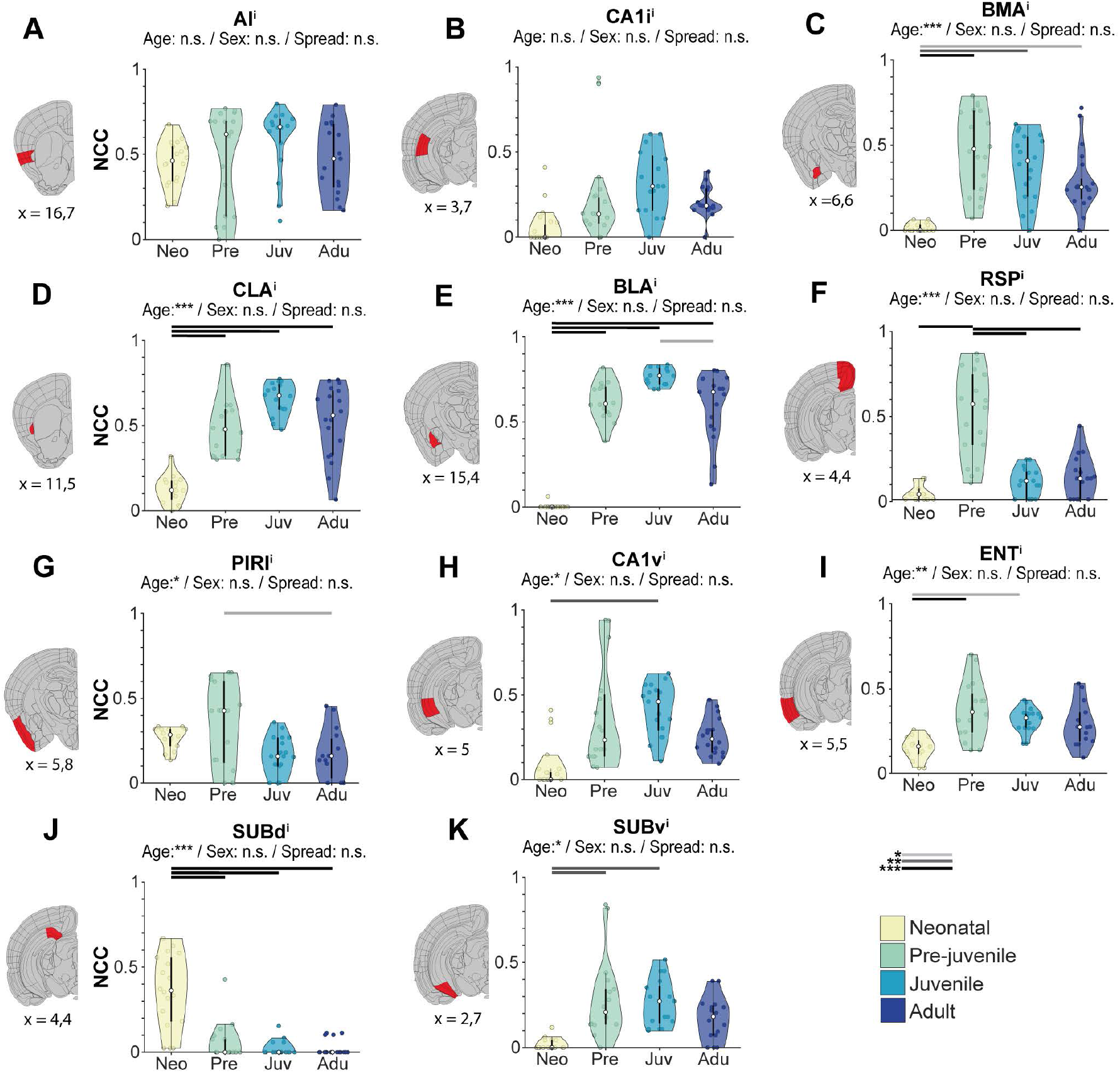

Thus, numerous ipsilateral projections target the mPFC during development and reveal distinct dynamics. Overall, long-range connectivity seems to be particularly dense during the pre-juvenile / juvenile period.

### 3.4. Prefrontal projections from thalamic and hypothalamic nuclei develop in an inverse manner

Next, we analyzed long-range prefrontal afferents arising from ipsilateral subcortical brain structures (ATN^i^, VM^i^, MD^i^, SUM^i^, PAG^i^, VTA^i^). For the ATN^i^, we quantified the interanterodorsal nucleus as well as ventral and dorsal sections of the AM. VTA^i^-to-mPFC projections remained unchanged throughout development (Figure 4A). In contrast, the density of VM^i^ and ATN^i^ afferents increased from neonatal to pre-juvenile age and remained stable afterwards, even though the density of projections at neonatal age was rather low in VM^i^ and high in ATN^i^ (Figure 4B,C). Notably, ATN^i^ was the only region for which we observed an effect of viral spread which decreased with age (see Supplementary discussion, Statistics table S1&S2). The density of labeled cells in the MD^i^ decreased with age, indicating a sparsification of prefrontal afferents (Figure 4D). Similarly, the density of SUM^i^-to-mPFC projections decreased from pre-juvenile to juvenile age with a significant effect of sex, revealing higher projection densities for male mice (Figure 4E, Statistics table S1&S2).

**Figure 4.**
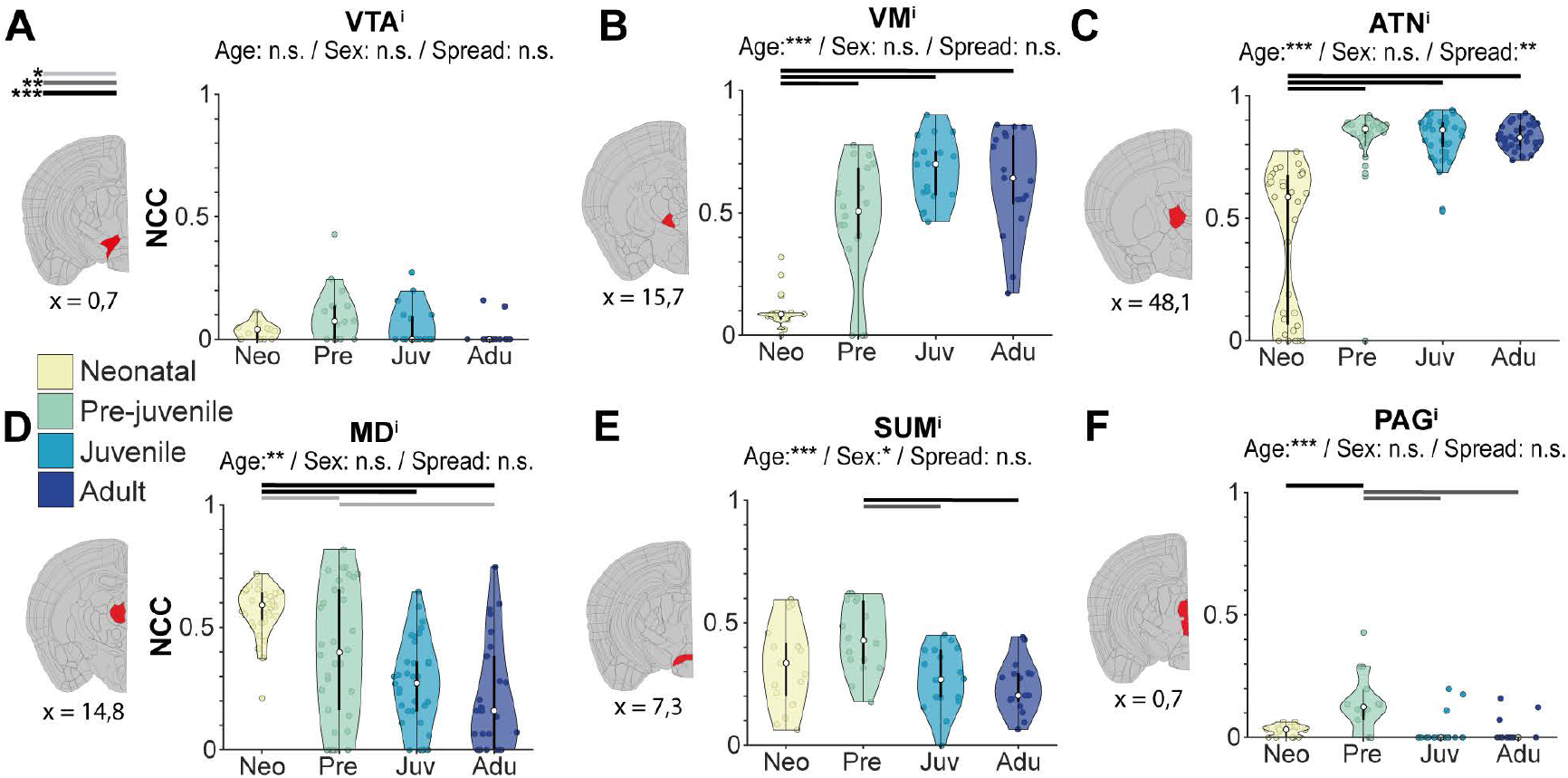

For the PAG^i^, the density of retrogradely labeled cells was generally low but followed a similar trend as most of the cortical areas and showed a peak at pre-juvenile age (Figure 4F).

Thus, most prefrontal afferents from subcortical regions reach high densities at pre-juvenile age, yet their dynamics, either increasing or decreasing with age, are region-specific.

### 3.5. Contralateral prefrontal afferents undergo pruning along development

Adult mPFC has been reported to receive afferents from numerous contralateral regions (marked as ^c^ for all regions) (Anastasiades and Carter, 2022). To clarify how these projections evolve along development, we monitored contralateral prefrontal afferents from birth to adulthood alongside ipsilateral projections. While no age-dependent changes of prefrontal afferent density from contralateral ACA^c^, ILA^c^ and MOs^c^ were detected (Figure S1A), the number of labeled cells in the PL^c^, similarly to the PL^i^, increased from neonatal to pre-juvenile age and remained stable afterwards (Figure 5A). In contrast, the number of labeled cells in the ORB^c^ decreased from neonatal to juvenile and adult age (Figure 5B). Whereas projection density from ORB^i^ showed no sex-dependent differences, an effect of sex was detected for the ORB^c^, the number of labeled cells being higher in male compared to female mice (see Statistics table S2). Other contralateral regions were affected neither by sex nor viral spread.

**Figure 5.**
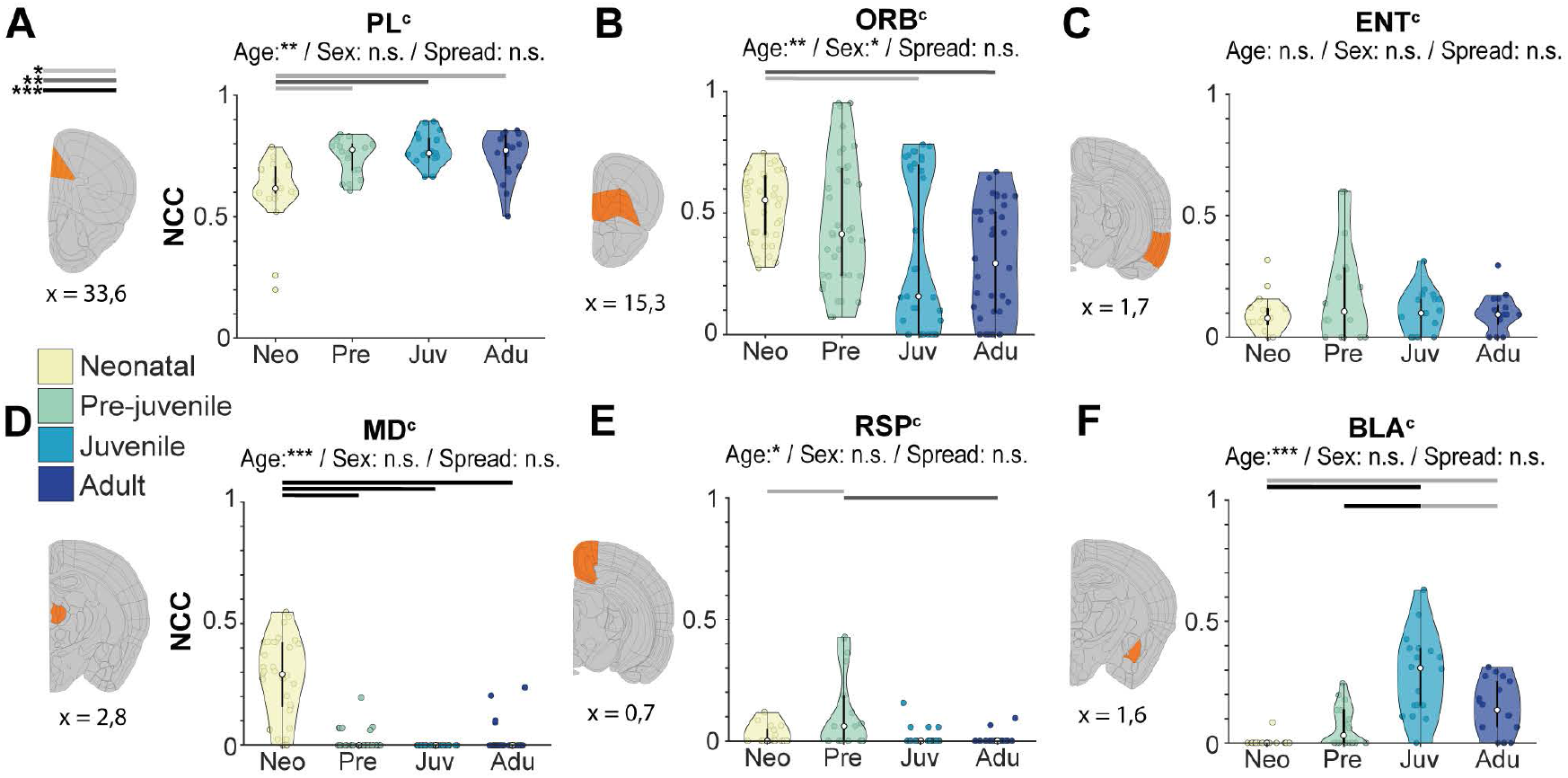

Based on their observed developmental dynamics, we classified contralateral long-range afferents into four distinct groups. First, afferents that were present at birth and their density remaining constant throughout development. The projections from ENT^c^ and CLA^c^ are characteristic for this pattern (Figure 5C; S1A). Second, afferents that were present early during development and strongly decrease later on. This patterns was detected for contralateral afferents from MD^c^, RSP^c^, AI^c^, ATN^c^, SUBd^c^, PIRI^c^, SUM^c^, and VTA^c^ (Figure 5D, E; S1B, C). The RSP^c^ was the only contralateral brain region for which we detected a density increase of retrogradely-labeled cells from neonatal to pre-juvenile age followed by a substantial decrease at older age (Figure 5E). Similar to the ORB^c^, VTA^c^ was one of the few regions that showed an age-dependent change in contra-but not ipsilateral projections to mPFC (Figure 5B; S1C). Third, the number of retrogradely labeled cells increased with age in a similar manner as their corresponding ipsilateral sections. Afferents from BLA^c^ and BMA^c^ belong to this group, despite the density differences between the neonatal and adult group being minor (Figure 5F; S1E). Fourth, almost no prefrontal afferents were detected during the entire development. This pattern was characteristic for CA1v^c^, CA1i^c^, SUBv^c^, VM^c^, and PAG^c^ (Figure S1D).

Taken together, these data reveal that the pruning of contralateral afferents along development follow age- and region-specific patterns, resulting in denser ipsilateral projections from most brain regions to the mPFC at adult age than those from the contralateral hemisphere.

### 3.6. Myelination of contralateral prefrontal afferents emerges after neonatal age

In addition to monitoring the density of contralateral prefrontal afferents, we also assessed their myelination. For this, we focused on the cortico-cortical projections crossing the CC.

The CC represents the largest white matter tract in the mammalian brain (Fitsiori et al., 2011; De León Reyes et al., 2020; Szczupak et al., 2021) and it has been extensively studied at adult age (Yorke Jr. and Caviness Jr., 1975; van der Knaap and van der Ham, 2011; De León Reyes et al., 2020; Innocenti et al., 2022). We characterized the developmental dynamics of myelination of contralateral prefrontal afferents in the previously defined age groups (neonatal, pre-juvenile, juvenile, and adult) (Figure 6A). Overall myelin expression increased with age (Figure 6B). Retrogradely labeled axons that colocalized with myelin were sparse in neonatal mice, but their density strongly increased at pre-juvenile age and remained elevated until adulthood (Figure 6C). Similarly, the portion of myelin colocalizing with labeled axons increased from neonatal to adult age (Figure 6D).

**Figure 6.**
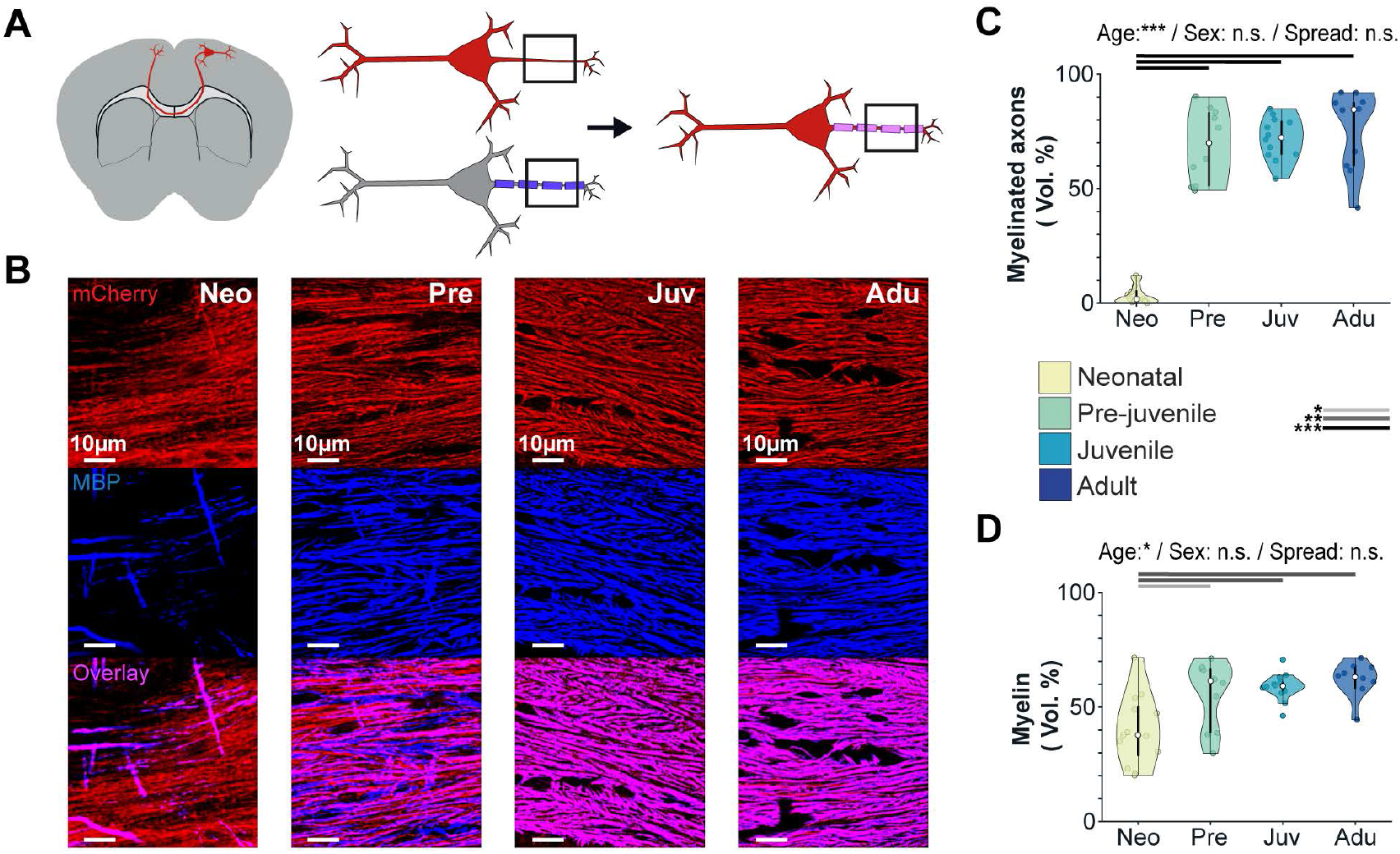

## 4. Discussion

The PFC is a higher-order cortical region that is essential for complex cognitive and affective processes such as decision making, working memory, and emotional regulation. Its maturation is protracted when compared to primary sensory cortices (Chini and Hanganu-Opatz, 2021), suggesting a prolonged time window for structural and functional refinement of prefrontal circuits. In this study, we unraveled the developmental dynamics of local and long-range ipsilateral and contralateral projections to the mPFC, with a specific focus on developmental changes in projection strength from different regions (*i.e*., ACA, PL, IL, ORB, MOs, AI, PIRI, BLA, BMA, RSP, ENT, CA1v, CA1i, SUBv, SUBd, CLA, ATN, VM, MD, SUM, PAG, and VTA) as well as the onset of myelination. Our findings reveal regionally distinct maturational dynamics of prefrontal afferents that might underlie functional and behavioral development.

### 4.1. Developmental shift from bilateral to unilateral projections to the mPFC

At birth, afferent projections to the mPFC are largely bilateral. However, this pattern shifts progressively towards ipsilateral projections during maturation, with only a subset of cortical and limbic areas retaining contralateral connectivity in adulthood (*i.e*., ACA^c^, PL^c^, ILA^c^, MOs^c^, ORB^c^, AI^c^, BLA^c^, BMA^c^, and ENT^c^). The most commonly observed dynamic of contralateral projections was a decline of projection density after neonatal age (*i.e*., ORB^c^, AI^c^, PIRI^c^, SUBd^c^, ATN^c^, MD^c^, SUM^c^). While postnatal pruning of perinatal contralateral projections for the ATN and MD has been reported (Minciacchi and Granato, 1989; De León Reyes et al., 2019), our results imply that this unilaterality of prefrontal projections might even be initiated before birth, as contralateral projections at neonatal age are already less dense than ipsilateral ones.

Our data show that myelination of CC-crossing prefrontal afferents begins only after the loss of most contralateral projections. In adult animals, the CC, is the largest white matter tract in the mammalian brain (Fitsiori et al., 2011; De León Reyes et al., 2020) and contains mostly cortico-cortical fibers (Szczupak et al., 2021), although evidence also suggests the presence of cortico-subcortical connections (Milardi et al., 2015; De Benedictis et al., 2016). In line with previous studies (Sturrock, 1980), we detected low amounts of myelin in the CC and found that mPFC axonal fibers are largely unwrapped at neonatal age. As has been previously shown (Paus et al., 1999; Fornari et al., 2007), myelination of prefrontal afferents increases strikingly during the second postnatal week, co-occurring with weaning and the onset of more complex behavioral abilities (Klune et al., 2021). As oligodendrocyte engagement and axonal wrapping are promoted by activity-dependent mechanisms (de Faria et al., 2019), the absence of myelin in early-stage interhemispheric fibers supports the idea that these projections might lack the necessary activity for functional embedding into mature circuits. In line with this hypothesis, the emergence of activity follows a protracted development in the mPFC (Brockmann et al., 2011). This transient bilateral connectivity may serve as a scaffolding mechanism for the maturation of local prefrontal microcircuits, facilitating early synchronization across hemispheres. The question of whether early contralateral prefrontal projections are functionally active remains to be addressed. Alternatively, these projections could reflect developmental remnants of cerebral ontogenesis that, in the absence of a specific function, are pruned with ongoing maturation.

In line with this, most ipsilaterally projecting regions show an overall increase of input strength from neonatal to adult age, some areas (*e.g*., MD^i^ and SUBd^i^) demonstrate an overall reduction in mPFC afferents, similar to their contralateral counterparts. This result partially contradicts earlier qualitative, not statistically validated reports, suggesting a rebound of MD input in adulthood (Rios and Villalobos, 2004). Given the well-established role of the MD in working memory and cognitive functions (Parnaudeau et al., 2018), developmental pruning of its projections to the mPFC appears paradoxical. However, pruning of excessive, non-informative, or redundant projections might improve signal fidelity. Our data suggest that this refinement of projections to the mPFC continues into adulthood, potentially facilitating the emergence of precise, goal-directed behavior.

### 4.2. Connectivity to the mPFC peaks during early adolescence

One of the most salient observations in our data is the pronounced increase of projections to the mPFC during early adolescence (*i.e*., pre-juvenile and juvenile group), coinciding with the emergence of higher-order cognitive functions (Klune et al., 2021; Thies et al., 2025). To this point, we could show that afferents from the limbic regions ENT^i^, SUBv^i^, CA1v^i^, BLA^i^, BLA^c^, and BMA^c^ peak in projection strength during early adolescence. For example, the SUBv is known to be involved in the reward circuit of the brain (Ding et al., 2020) and a role of the CA1 in anticipation of consequences has been proposed (Fyhn et al., 2002; Numan, 2015). Thus, this temporal alignment of projections to the mPFC supports behavioral studies that implicate adolescence as a sensitive window for risk taking and cognitive flexibility in humans (van Duijvenvoorde et al., 2016; McCormick and Telzer, 2018) and in rodents (Johnson and Wilbrecht, 2011; Klune et al., 2021; Thies et al., 2025). Similarly, projections from the amygdala (*i.e*., BLA and BMA of both hemispheres) are nearly absent in neonatal mice but strongly increase towards pre-juvenile age. This developmental trajectory has been previously described (Cunningham et al., 2002) and might be essential for the consolidation of fear memories. Notably, the BLA has also been shown to play a role in decision-making paradigms, in which effort and reward values have to be evaluated (Floresco and Ghods-Sharifi, 2007). Moreover, increased projection strength from the AM (as part of the ATN^i^) to the mPFC could facilitate dopamine-dependent reward processing, as its role in reinforcing goal-directed behaviors has been shown in adult mice (Yang et al., 2022). Since the density of dopaminergic receptors in the mPFC peaks during adolescence (Larsen and Luna, 2018), the maturation of these inputs might contribute to the anatomical foundation for adaptive decision-making and behavioral flexibility.

While some brain areas maintained their ipsilateral prefrontal innervations strength after initial increase, other regions showed a decline of projections after early adolescence (*i.e*., PIRI^i^, RSP^i^, and SUM^i^). These brain regions are associated with olfactory processing, spatial navigation, and episodic memory, all of which become particularly relevant during the transition from maternal dependency to independent exploration (Vann et al., 2009; Ito et al., 2018; Loureiro et al., 2019). The PIRI projects to the ORB as well as to the mPFC (Ährlund-Richter et al., 2019; Bhattarai et al., 2022). These pathways have been reported to be relevant for assessment of food safety (Loureiro et al., 2019). The SUM and RSP are involved in both prefrontal and hippocampal networks and play a role in spatial navigation as well as episodic memory (Vann et al., 2009; Ito et al., 2018). We hypothesize that transient innervation of the early adolescent mPFC by these areas facilitates critical behavioral adaptations at the time of weaning, including food selection and environmental orientation.

### 4.3. Concluding remarks

In summary, our data reveal that the development of mPFC afferents is neither linear nor uniform but characterized by region-specific dynamics of input formation, stabilization, and pruning (see Supplementary discussion). These anatomical changes closely parallel known milestones of behavioral and functional maturation, suggesting structural development as a framework for functional circuit integration and adult-like cognitive functions. These findings help to understand how early afferent maturation shapes the integrative functions of the mPFC and how disruptions may result in lifelong lasting phenotypes, such as those associated with neurodevelopmental disorders.

## 6. CRediT authorship contribution

Conceptualization, supervision, and project administration: JAP, AG, and ILH-O. Investigation: MS, AG, JAP, AF, and AM. Methodology and validation: AG and JAP. Data curation, formal analysis, and visualization: MS, JAP, and AG. Writing – original draft: JAP, AG, and MS. ILH-O, JAP, AG, and TO: Writing – review and editing. All authors have approved the final version of the manuscript.

## 8. Acknowledgments

We thank P. Putthoff and A. Dahlmann for excellent technical assistance.

## 9. Resource availability

### Lead contact

Jastyn A. Pöpplau

### Materials availability

The study did not generate new unique reagents.

### Data and code availability

All data needed to evaluate the conclusions are available in the figures, main text, or the supplementary materials. Raw images are available from the lead contact upon request. Custom image analysis code is available from the lead contact upon request.

## 10. Glossary

mPFC: medial prefrontal cortex
ACA: anterior cingulate cortex
PL: prelimbic cortex
IL: infralimbic cortex
ORB: orbitofrontal cortex
MO: secondary motor cortex
AI: agranular insular cortex
PIRI: piriform cortex (),
BLA: basolateral amygdala,
BMA: basomedial amygdala
RSP: retrosplenial cortex
ENT: entorhinal cortex
CA1v: ventral CA1
CA1i: intermediate CA1
SUBv: ventral subiculum
SUBd: dorsal subiculum
CLA: claustrum
ATN: anterior nuclei
VM: ventromedial thalamus
MD: mediodorsal thalamus
SUM: supramammillary nucleus
PAG: periaqueductal gray
VTA: ventral tegmental area
NCC: normalized cell count
rAAV: recombinant adeno-associated virus
P: postnatal day

