## Supplementary Information for "Mapping prefrontal afferents along development"

**Inventory of Supplementary Information**


| **Inventory of Supplementary Information** | Page |
|---|---|
| Supplementary discussion | 3 |
| Figure S1 | 5 |

### SUPPLEMENTARY DISCUSSION

#### Local mPFC connectivity

Local prefrontal circuits rely on input from long-range projections in order to process incoming information and execute prefrontal functions (Hanganu-Opatz et al., 2023). We observed slight regional specificity of mPFC projection dynamics. While prefrontal projection strength from PL$^i$ and ILA$^i$ increased from neonatal to adult age, the ACA$^i$ showed an increase towards pre-juvenile age but no difference between neonatal and adult age. Notably, the PL$^c$ was the only prefrontal subdivision that showed an increase in prefrontal projection strength from neonatal to adult age. In contrast, the ORB$^c$ showed a decline in contralateral projections. These divergent trajectories potentially reflect distinct cytoarchitectural differentiation between prefrontal areas during maturation of callosal connectivity (Barbas et al., 2005; Kolk and Rakic, 2022), possibly reflecting their contributions to cognitive, emotional, and sensorimotor processing. On a functional level, previous work identified a peak in broadband gamma and spiking activity in the mPFC during early adolescence which orchestrates a functional reorganization of mPFC circuits. This period of heightened activity is followed by a reduction, ultimately leading to mature mPFC synchrony at adult age (Pöpplau et al., 2024). Thus, the temporal rise in external limbic afferents may serve as instructive signals for the refinement of mPFC microcircuits.

#### Methodological Considerations

The retrograde labeling virus rAAV9rg_CaMKII-mCherry does not exclusively infect excitatory neurons via axonal pathways but can also infect neurons in close proximity via their somata. Therefore, quantification of labeled cells, especially in subdivisions of the ipsilateral mPFC could have been biased by an uneven distribution of the retrograde labeling virus. Furthermore, the only identified increase of interhemispheric projection strength, was observed in the PL$^c$, which could possibly be explained by a bias due to experimental procedures. Homotopic regions are more strongly connected to each other than heterotopic ones (Barbas et al., 2005) and as the PL is located between the ACA and ILA on the dorso-ventral axis, it is most likely that the PL$^i$ received the largest portion of viral vector during the injection procedures resulting in the PL$^c$ being labeled more strongly than the heterotopic ACA$^c$ and ILA$^c$.

The ATN$^i$ was the only brain region influenced by the virus spread which might be due to its composition as a group of several thalamic nuclei. Individual anterior thalamic nuclei project to the mPFC, but also adjacent brain regions (Aggleton and O'Mara, 2022). Thus, it is possible that a higher virus spread into adjacent brain regions leads to higher cell counts in the ATN. Especially viral spread into the ORB which has been shown to receive direct input from the AM (Carmichael and Price, 1995), might have contributed to a significant effect of viral spread.

**Notes on Specific Brain Areas**

We included the quantification of very sparsely labeled cells from the midbrain in our analysis, which have not been reported to project to the adult mPFC via excitatory glutamatergic fibers in previous studies. Prior research shows that the PAG receives projections from the mPFC (Anastasiades and Carter, 2021) but no study provides evidence for reciprocal connectivity. The VTA is known to innervate the mPFC via dopaminergic connections (Tzschentke, 2000), but glutamatergic projections from the VTA to the mPFC have not been described before. The absence of reports of glutamatergic prefrontal afferents from the PAG and VTA may result from the very low number of prefrontally projecting cells and/or the transient nature of these projections during development, as our results suggest. Finally, in our analysis we could not detect any projections from the SUBd to the mPFC after neonatal age, while a previous study described prefrontal afferents from the SUBd in adult animals (Strange et al., 2014) but without identifying such projections consistently for all experimental subjects. The cause of such conflicting results remains unclear and requires further experimental research.

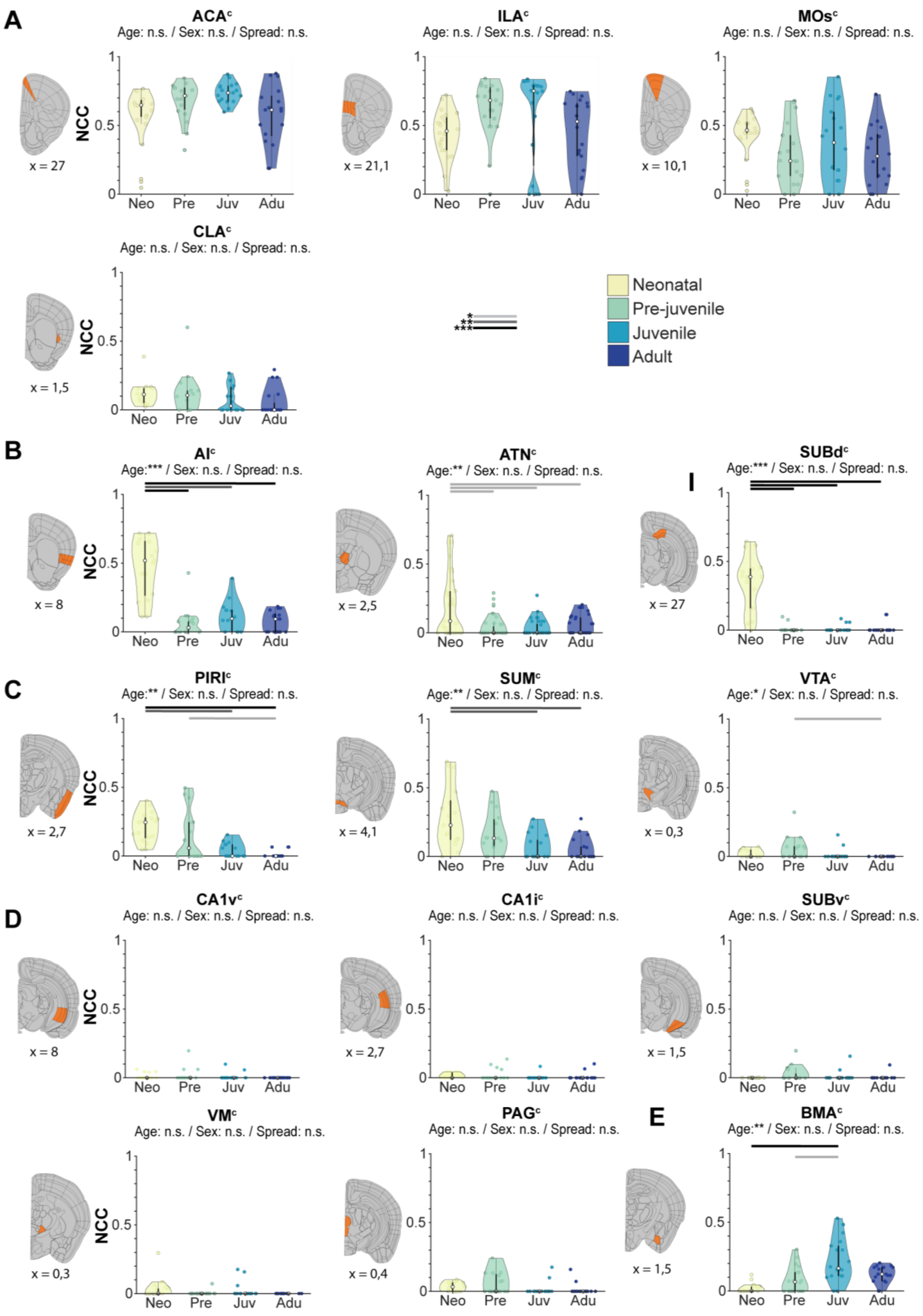
A
ACA^c
Age: n.s. / Sex: n.s. / Spread: n.s.
ILA^c
Age: n.s. / Sex: n.s. / Spread: n.s.
MOs^c
Age: n.s. / Sex: n.s. / Spread: n.s.
CLA^c
Age: n.s. / Sex: n.s. / Spread: n.s.
NCC
Neo Pre Juv Adu
x = 27
x = 21,1
x = 10,1
x = 1,5
Neonatal
Pre-juvenile
Juvenile
Adult
*
**
***
B
AI^c
Age:*** / Sex: n.s. / Spread: n.s.
ATN^c
Age:** / Sex: n.s. / Spread: n.s.
SUBd^c
Age:*** / Sex: n.s. / Spread: n.s.
x = 8
x = 2,5
x = 27
C
PIRI^c
Age:** / Sex: n.s. / Spread: n.s.
SUM^c
Age:** / Sex: n.s. / Spread: n.s.
VTA^c
Age:* / Sex: n.s. / Spread: n.s.
x = 2,7
x = 4,1
x = 0,3
D
CA1v^c
Age: n.s. / Sex: n.s. / Spread: n.s.
CA1i^c
Age: n.s. / Sex: n.s. / Spread: n.s.
SUBv^c
Age: n.s. / Sex: n.s. / Spread: n.s.
x = 8
x = 2,7
x = 1,5
VM^c
Age: n.s. / Sex: n.s. / Spread: n.s.
PAG^c
Age: n.s. / Sex: n.s. / Spread: n.s.
E
BMA^c
Age:** / Sex: n.s. / Spread: n.s.
x = 0,3
x = 0,4
x = 1,5

**FIGURE S1 related to FIGURE 5 | Age-dependent dynamics of contralateral projections to the mPFC**. **(A)** Top left, Violin plots displaying the NCC of prefrontally projecting $ACA^c$ cells (n = 72 images, 23 mice) for each age group. Top middle, same as top left for $ILA^c$ (n = 80 images, 24 mice). Top right, same as top left for $MOs^c$ (n = 70 images, 24 mice). Bottom left, same as top left $CLA^c$ (n = 69 images, 23 mice). **(B)** Left, same as (A) for $AI^c$ (n = 64 images, 24 mice). Middle, same as (A) for $ATN^c$ (n = 135 images, 24 mice). Right, same as (A) for $SUBd^c$ (n = 68 images, 24 mice). **(C)** Left, same as (A) for $PIRI^c$ (n = 63 images, 22 mice). Middle, same as (A) for $SUM^c$ (n = 65 images, 24 mice). Right, same as (A) for $VTA^c$ (n = 72 images, 23 mice). **(D)** Top left, same as (A) for $CA1v^c$ (n = 132 images, 24 mice). Top middle, same as (A) for $CA1i^c$ (n = 65 images, 24 mice). Top right, same as (A) for $SUBv^c$ (n = 65 images, 24 mice). Bottom left, same as (A) for $VM^c$ (n = 69 images, 24 mice). Bottom middle, same as (A) for $PAG^c$ (n = 67 images, 24 mice). **E)** Same as (A) for $BMA^c$ (n = 71 images, 24 mice). Assessed contralateral areas are indicated in orange in schematics on the left with mean cell count given as x below sections. Violin plots are represented as median with $25^{th}$ to $75^{th}$ percentile. See Statistics table S1 for details.
